# The two plant-specific DREAM components FLIC and FLAC repress floral transition in *Arabidopsis*

**DOI:** 10.1101/2023.08.07.552284

**Authors:** Lucas Lang, Franziska Böwer, Hasibe Tunçay Elbaşı, Dominique Eeckhout, Nick Marschlich, Geert de Jaeger, Maren Heese, Arp Schnittger

## Abstract

The DREAM complex is a key transcriptional regulator especially involved in the control of the cell cycle and development. Here, we characterise two novel plant- specific DREAM components, FLIC and FLAC, which we identified through tandem affinity purification experiments as interactors of conserved core DREAM constituents. We demonstrate that plants lacking both FLIC and FLAC exhibit pleiotropic phenotypes, including stunted growth and reduced fertility. Notably, *flic flac* double mutants show an early-flowering phenotype, an aspect that we found to be shared with mutants of the core DREAM component LIN37, with which FLIC and FLAC interact in binary protein-protein interaction assays. Performing reverse affinity purification experiments, we detected the JMJ14/NAC050/NAC052 module, known for its involvement in flowering repression, in the interactome of both FLIC and FLAC. Subsequent binary interaction studies then link the JMJ14/NAC050/NAC052 module via LIN37 to the DREAM complex providing a mechanistic framework on how flowering time could be transcriptionally controlled by the DREAM complex.

**Summary blurb:** This study identifies two plant-specific members of the DREAM complex, explores their roles by mutant analysis and protein interaction investigation, and links them and additional DREAM complex components to the regulation of floral transition.

## Introduction

Proper timing of the transition from vegetative growth to flowering is of crucial importance for a plant to maximise its reproductive success. Reflecting the importance and complexity of this transition, an intricate genetic network gauges both the endogenous state of a plant as well as the environmental conditions and promotes flowering at the optimal moment (Koornneef et al, 1998; Amasino & Michaels, 2010; Srikanth & Schmid, 2011).

During the extensive research of flowering time genes that has been performed in Arabidopsis, two major networks have been identified (Simpson & Dean, 2002; Kinoshita & Richter, 2020). First, internal factors, such as plant age and phytohormones, regulate initiation of flowering (partly) independently of environmental cues (Huijser & Schmid, 2011; Hyun et al, 2017). Secondly, floral transition can be controlled externally by factors such as vernalisation, photoperiod, and temperature (Yanovsky & Kay, 2002; Andrés & Coupland, 2012).

Many floral effector signals become integrated at the *FLOWERING LOCUS T* (*FT*) gene which is required for induction of floral meristem identity (Wigge et al, 2005; Hayama et al, 2017). To initiate floral transition, FT gets relocated to the shoot apical meristem by florigen transporters (Abe et al, 2005; Corbesier et al, 2007; Jaeger & Wigge, 2007), where it then forms the florigen activation complex (FAC) to regulate floral meristem identity genes, such as *APETALA1* (*AP1*), *FRUITFULL* (*FUL*), *SUPPRESSOR OF OVEREXPRESSION OF CONSTANS1* (*SOC1*), and *SEPALLATA3* (*SEP3*) (Abe et al, 2005; Kawamoto et al, 2015; Collani et al, 2019).

*FT* expression is also subject to epigenetic regulation. Several Jumonji C (JmjC) domain-containing histone demethylases have been demonstrated to regulate *FT* in different ways. The H3K27 demethylase RELATIVE OF EARLY FLOWERING 6 (REF6)/JUMONJI 12 (JMJ12) activates *FT* transcription when overexpressed, whereas its close homologue EARLY FLOWERING 6 (ELF6) is an upstream repressor of photoperiodic floral induction (Noh et al, 2004; Lu et al, 2011). Additionally, there are H3K4-specific members of the JMJ demethylase group that either repress *FT*, such as JMJ14, or activate *FT*, such as JMJ15 and JMJ18 (Lu et al, 2010; Yang et al, 2012a, 2012b). One central regulator of florigen expression is the well-described MADS-box transcription factor FLOWERING LOCUS C (FLC) (Wang et al, 2014). Recently, it has been demonstrated that the floral repression through FLC can be counteracted by absence of JMJ14 through increased H3K4me3 levels, emphasising its role as an important epigenetic flowering time regulator (Richter et al, 2019). JMJ14 physically associates with the plant-specific NAM, ATAF1, and CUC1/CUC2 (NAC) type transcription factors NAC050 and NAC052, and it has been demonstrated that a similar set of genes is deregulated in *jmj14* and *nac050 nac052* mutants leading to an early-flowering phenotype in both mutants (Ning et al, 2015; Zhang et al, 2015). Further, an interaction of JMJ14 with TELOMERE REPEAT BINDING FACTOR (TRB) proteins has been shown recently, together with hyper-methylation of a common set of genes in the respective mutants as well as an impact on flowering time regulation via the naturally silenced transcription factor FLOWERING WAGENINGEN (FWA) (Soppe et al, 2000; Wang et al, 2023).

Developmental transitions, such as the induction of flowering, also usually involve a fine-tuned cell proliferation control (Jacqmard et al, 2003). The DP, RB-like, E2F, and multi-vulval class B (MuvB)-core (DREAM) complex is a multi-protein complex with a canonical role in transcriptional cell cycle regulation (Korenjak et al, 2004; Litovchick et al, 2007; Schmit et al, 2007; Asthana et al, 2022; Koliopoulos et al, 2022). Notably, mutants in several components of the DREAM complex have been previously linked to control of floral transition (Bouveret et al, 2006; Ning et al, 2020). However, this complex has only recently been fully identified in plants and how this complex and/or subcomplexes control flowering is largely not understood (Ning et al, 2020; Lang et al, 2021).

The MuvB-core is considered to be the constant part that is present in every variation of the DREAM complex and is comprised of the different homologues of ALWAYS EARLY (ALY, a homologue of the *C. elegans* DREAM component LIN9), ABNORMAL CELL LINEAGE PROTEIN 37 (LIN37), LIN52, TESMIN/TSO1-LIKE CXC (TCX, a LIN54 homologue), and MULTICOPY SUPPRESSOR OF IRA (MSI, a RBBP4 homologue). This core complex is able to recruit other proteins, such as RETINOBLASTOMA-RELATED 1 (RBR1), DIMERIZATION PARTNER (DP), E2 PROMOTER BINDING FACTOR (E2F), and MYELOBLASTOMA 3R (MYB3R), enabling the DREAM complex to control and maintain the appropriate transcriptional program (Kobayashi et al, 2015; Fischer & Müller, 2017). It has been postulated that depending on the exact homologues being incorporated in a certain variant of the DREAM complex, it can facilitate either activation or repression (or both for different subsets of genes) (Magyar et al, 2016; Fischer & Müller, 2017).

Recent research on the plant DREAM complex has focused on DNA methylation maintenance, developmental control, and the DNA damage stress response (Ning et al, 2020; Lang et al, 2021; Wang et al, 2022b). Late-flowering phenotypes have been described for mutants of the MuvB-core components *MSI1*, which also constitutes a subunit of the Polycomb Repressive Complex 2 (PRC2), as well as for *TCX5* and *TCX6* (Bouveret et al, 2006; Ning et al, 2020). MSI1 is responsible for gene silencing by controlling histone methylation and acetylation levels, and mutation leads to reduced expression of CO. This results in the inability of proper *FT* and *SOC1* activation, thereby inhibiting the photoperiod pathway of floral initiation (Steinbach & Hennig, 2014). Furthermore, it has recently been shown that MSI1 is needed interdependently with HISTONE DEACETYLASE 6 (HDA6) for repression of the flowering repressor genes *FLC*, *MADS AFFECTING FLOWERING 4* (*MAF4*), and *MAF5* (Xu et al, 2022). Here, we have identified a homologous pair of uncharacterised, plant-specific interactors of the DREAM complex, FLORAL INDUCTION CONTROLLER (FLIC) and FLORAL ANTICIPATION CONTROLLER (FLAC). By performing different interaction assays, we show how they integrate into the complex, and we characterise them as novel DREAM components in plants. Observing the phenotypes of the *flic flac* double mutant, we describe its pleiotropic effects, ranging from severe vegetative defects to strongly affected fertility, indicating a role in a wide range of developmental processes. We further demonstrate a photoperiodically independent early floral transition in the absence of FLIC and FLAC as well as for mutants of the MuvB-core component LIN37 and connect them to the JMJ14/NAC050/NAC052 module.

## Results

### FLIC and FLAC are novel components of the plant DREAM complex

Previously, we have identified several uncharacterised proteins as potential DREAM complex interactors by tandem affinity purification (TAP) analyses conducted with TCX5, LIN52, and RBR1 serving as baits (Lang et al, 2021). Amongst those, AT5G10110, which we have here named FLORAL INDUCTION CONTROLLER (FLIC), co-purified as a prey of TCX5. A similarity search using BLAST identified no homologues outside of the plant kingdom, while one homologous protein, AT5G65120 (FLORAL ANTICIPATION CONTROLLER, FLAC), was identified in *Arabidopsis thaliana*. Aligning the sequences of FLIC and FLAC reveals a sequence identity of 47.2% (Fig S1), indicating an overlapping function (Pearson, 2013).

To explore their interaction network, we first performed reciprocal affinity purification coupled to mass spectrometry (AP-MS) analyses using N- and C-terminal fusions of these two proteins as baits. For each experiment, three replicates were conducted. A total of 37 interacting proteins were identified in at least two replicates (Tables S1 and S2). Notably, homologues of the entire MuvB-core were found (Fig 1, Tables S1 and S2). Furthermore, the atypical E2F transcription factors E2FE/DEL1 and E2FF/DEL3 could be retrieved with both FLIC and FLAC baits expanding the previous finding that E2FE/DEL1 can be co-purified with the DREAM component TCX5 (Lang et al, 2021). Interestingly, we also saw the co-purification of the recently described, plant-specific transcriptional repressors BARRIER OF TRANSCRIPTIONAL ELONGATION 1 (BTE1)/DREAM COMPONENT 2 (DRC2) and BTE1-LIKE 1 (BTL1) which have been previously identified as DREAM interactors (Derkacheva et al, 2013; Ning et al, 2020; Lang et al, 2021; Wang et al, 2022b). A single MYELOBLASTOMA 3R (MYB3R) transcription factor could be detected, i.e. MYB3R1 appeared as a prey of FLIC.

**Figure 1.**
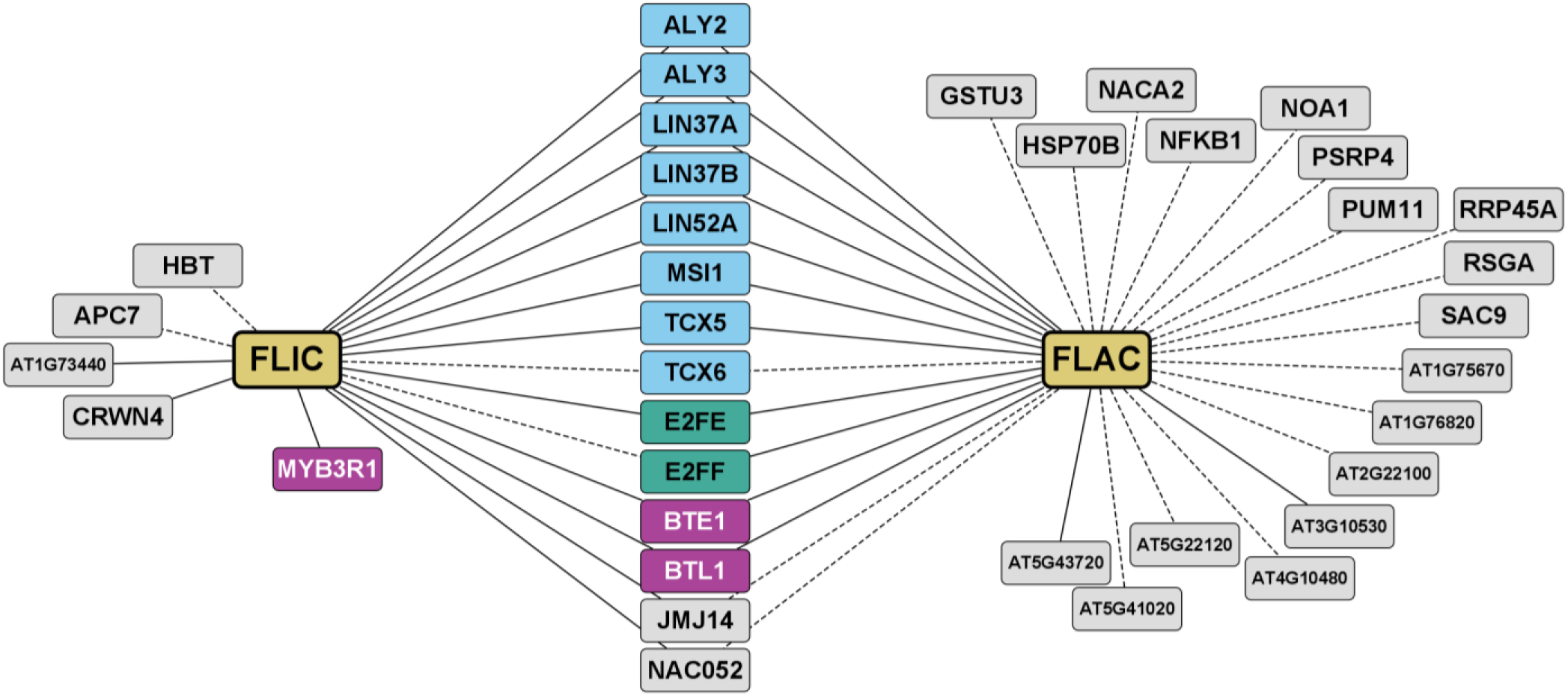
Overview of AP-MS results using FLIC and FLAC as baits. FLIC and FLAC were used as baits, N- as well as C-terminally tagged, in three replicates per fusion. All prey proteins that were found in at least two experiments and passed the filtering thresholds specified in the method section are connected to the respective bait by an edge. Solid edges represent prey found with both N- and C-terminally tagged bait, while prey connected by dashed edges were only detected above the threshold with either tag position. Gold, FLIC/FLAC; light blue, MuvB-core protein; dark cyan, other DREAM protein; pink, DREAM-related proteins; grey, other proteins.

Interestingly, amongst a group of 24 proteins that have so far not been linked to the DREAM complex, we noted the presence of the histone demethylase JUMONJI 14 (JMJ14) as well as the NAC type transcription factor NAC052 as preys for both FLIC and FLAC. JMJ14, NAC052, and its close homologue NAC050 have been shown to form a complex that is involved in flowering time regulation (Ning et al, 2015; Zhang et al, 2015; Rodrigues et al, 2021). Other hits include proteins involved in various processes, e.g. translational control and protein degradation. This suggests that FLIC and FLAC might be relevant in several additional pathways that may or may not be connected to DREAM function.

### FLIC and FLAC bind to the MuvB-core

To further investigate the relationship between FLIC, FLAC, and the Arabidopsis DREAM complex, we investigated binary interactions by performing yeast two-hybrid (Y2H) assays, including BTE1 and BTL1, that have been identified in previous TAP experiments. Noticeably, FLIC exhibited a strong auto-activational activity when fused to the DNA-binding domain (BD), so that interactions could not be checked with the BD-FLIC fusion protein.

FLIC and FLAC both interacted with LIN37, LIN52, BTE1, and MYB3R3 (Fig 2A and B). This finding also confirms an interaction between FLIC and BTE1 that has been shown before (Arabidopsis Interactome Mapping Consortium, 2011). Whereas FLIC showed no interactions specific to a certain homologue and interacted with all of LIN37A, LIN37B, LIN52A, LIN52B, BTE1, and BTL1, FLAC appears to have a more defined interaction activity with a single homologue of each protein group, i.e. only with LIN37B, LIN52A, and BTE1. Additionally, we could see BD-FLAC binding to CDKA;1, which has previously been suggested to be part of plant DREAM complexes (Kobayashi et al, 2015; Lang et al, 2021). Although both FLIC and FLAC showed interactions with MYB3R3, which co-purified with TCX5 as a TAP bait, interactions could not be observed with MYB3R1, which we identified in DREAM-related TAP experiments before (Lang et al, 2021).

**Figure 2.**
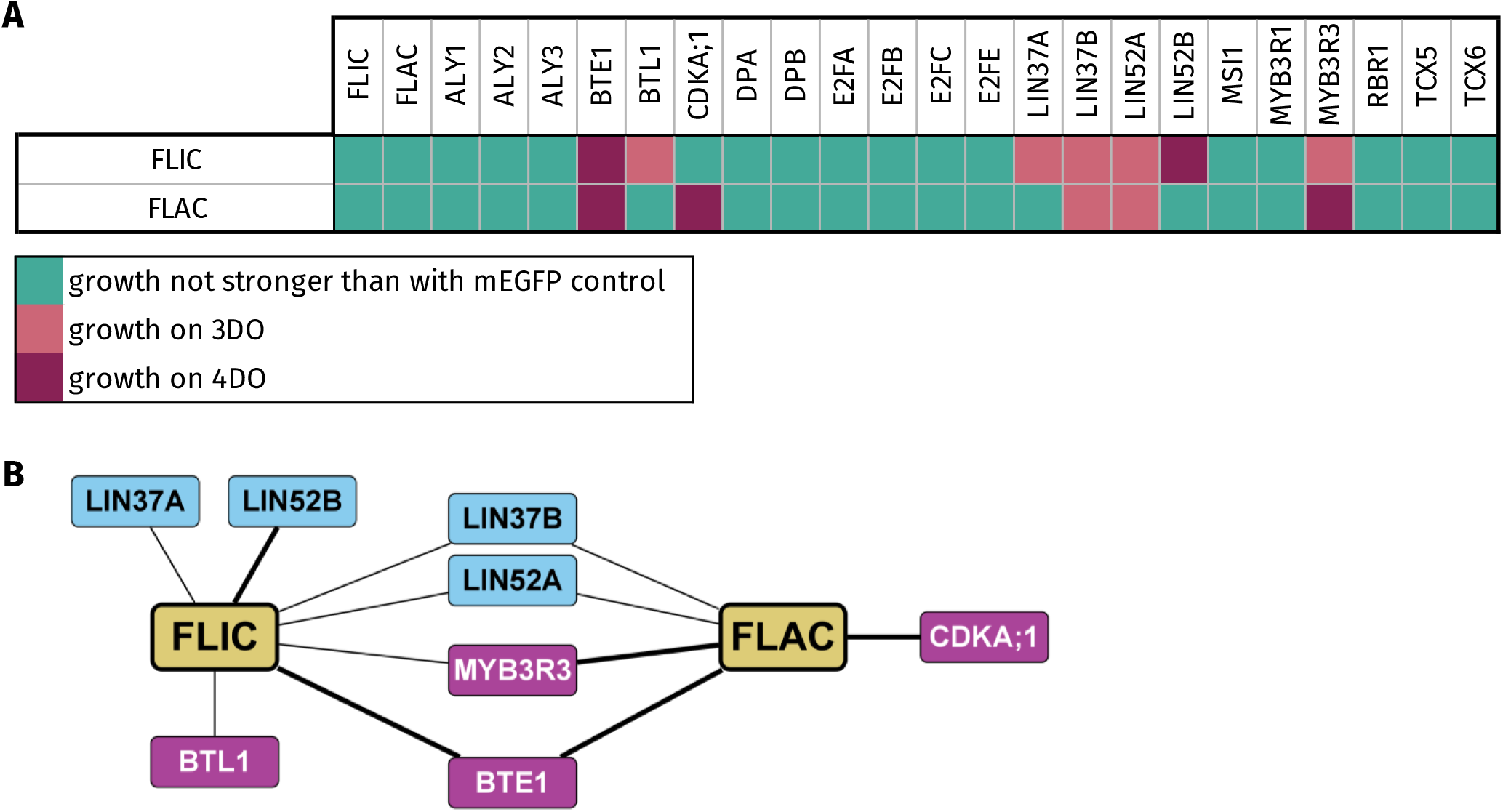
Binary interactions of FLIC and FLAC with the DREAM complex and associated proteins as determined by Y2H. Combinations for FLIC were only tested using the AD-FLIC fusion, as BD-FLIC showed strong auto-activation as observed with an AD-mEGFP control. FLAC was tested using both variants and only the strongest interaction is shown. Signal strength was classified according to yeast growth on different dropout media in three categories. **(A)** Dark pink, growth on 4DO; red, growth on 3DO but not on 4DO; cyan: growth not stronger than with mEGFP control. **(B)** Schematic representation of the observed interactions of FLIC and FLAC indicated by edges. Thick line, growth on 4DO; thin line, growth on 3DO but not on 4DO. Gold, FLIC/FLAC; blue, MuvB-core protein; pink, DREAM-related proteins.

Since we could see strong interactions of FLIC and FLAC with BTE1 in the Y2H, as well as with both BTE1 and BTL1 in the AP-MS experiments (Fig 2A and B, Table S1), we also investigated how BTE1 and BTL1 bind to the DREAM complex (Fig S2) and found that they appear to only connect to the MuvB-core via LIN37B. Apart from that, both BTE1 and BTL1 show an interaction with MYB3R3. Additionally, BTE1 appears to be binding DPA and E2FE, whereas BTL1 binds MYB3R1.

Taken together, these results suggest that FLIC and FLAC can act as part of a plant DREAM complex and bind to the MuvB-core via a specific subset of its components, namely LIN37 and LIN52.

### Concomitant loss of *FLIC* and *FLAC* leads to defects in vegetative growth and reduced fertility

To address the function of *FLIC* and *FLAC*, we characterised T-DNA insertion mutants for both genes. One transcriptional null mutant for each gene could be identified, *flic*-1 and *flac*-1 (Fig S3), and we refer to them in the following simply as *flic* and *flac*. Since both single mutants did not exhibit any striking mutant phenotype by themselves, we generated a *flic flac* double mutant. Notably, we only detected 15% double homozygous mutants (N = 184) with an excess of wild-type plants (34%) when the progeny *flic^-/-^ flac^+/-^* was analysed.

Homozygous double mutant plants exhibit several differences in vegetative growth when compared to the wildtype and the single mutants (Fig 3A and B). First, rosette leaf size is generally reduced and a serrated leaf shape is observable. Next, flowering starts early with the formation of multiple shoots that are thinner than in the wildtype, indicating a loss of apical dominance. This results in a bushy appearance of adult plants with reduced stem height. Furthermore, silique size and pollen amount are drastically reduced in the double mutant.

**Figure 3.**
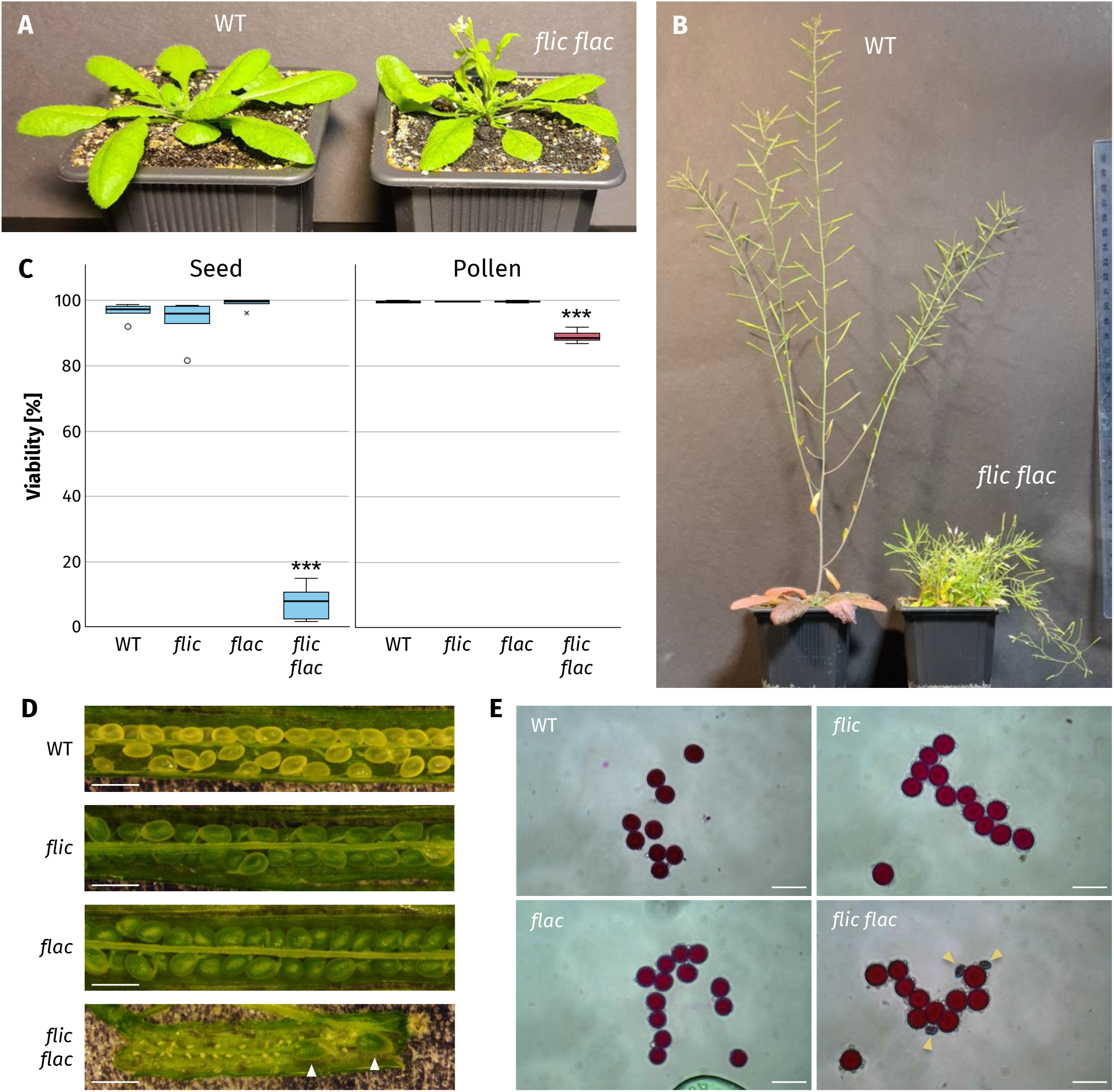
Phenotypical analysis of *flic flac* mutant plants. **(A)** Wildtype (WT, left) and *flic flac* mutant (right) grown under long-day conditions. An early-bolting phenotype can be observed in *flic flac* mutants. **(B)** Adult wildtype (left) and *flic flac* mutant (right) grown under long-day conditions. The *flic flac* double mutants show a loss of apical dominance resulting in a bushy phenotype and dramatically reduced height. **(C)** Quantification of seed and pollen viability in wildtype, *flic*, *flac*, and *flic flac* mutants. Seed viability analysis was conducted with 5 siliques from 5 plants per genotype. For assessing pollen viability, 9 flowers were analysed for wildtype, *flic*, and *flac* mutants, and 15 flowers were analysed for *flic flac* mutants. **(D)** Representative open siliques of wildtype, *flic*, *flac*, and *flic flac* mutants. White arrowheads show viable seeds in the *flic flac* silique. Scale bars, 1 mm. **(E)** Representative Peterson staining of wildtype, *flic*, *flac*, and *flic flac* pollen. Golden arrowheads show aborted pollen grains. Scale bars, 100 µm.

An analysis of seed viability revealed that only 7.5±2.5% (*P* < 0.001) of *flic flac* seeds are viable, whereas there was no significant reduction of viability in the single mutants (*flic*: 93.5±3.1%; *flac*: 99.0±0.7%) in comparison with the wildtype (96.5±1.2%) (Fig 3C and D). This reduction is consistent with the above-described strong reduction of viable seeds in double mutant siliques.

To test whether FLIC and FLAC are also required for gametophyte development, we performed reciprocal crossing between wild-type and *flic flac*^+/−^ plants and quantified the proportion of *FLAC*^+/−^ progeny. We could detect that loss of *FLAC* in either of the gametophytes resulted in a reduced number of heterozygous *FLAC* F1 plants (34% and 36%, respectively; N = 152–183), demonstrating that at least one functional copy of *FLIC* or *FLAC* is required for faithful female and male gamete transmission. Consistent with a requirement of FLIC and FLAC for male gametophyte development, we found that pollen viability in the double mutant is reduced (Fig 3C and E). Whereas viable pollen grains made up 99.6±0.2% of all pollen in the wildtype (*flic*: 99.7±0.1%; *flac*: 99.8±0.2%), this proportion is reduced to 89.1±0.9% in *flic flac* (*P* < 0.001).

These mutant phenotypes indicate that FLIC and FLAC are redundantly required for various aspects of sporophyte and gametophyte development.

### FLIC and FLAC control flowering time

As indicated by our AP-MS analyses, FLIC and FLAC appeared to be possible interactors of the floral transition regulators JMJ14 and NAC052. Since early flowering was one of the strongest and most obvious mutant phenotypes of the *flic flac* double mutant, we focussed next on the description of this defect. We first compared the bolting time as well as the number of rosette leaves at bolting of the *flic* and *flac* single mutants as well as the *flic flac* double mutant under long-day (LD) conditions in a 16 h light/8 h dark cycle. For this purpose, days after sowing (das) were tracked and leaves were counted as soon as the shoot of a plant reached a length of 1 cm.

Our results show that whereas the *flic* single mutant (31.2±0.5 das; 19.0±0.9 leaves) does not differ significantly from the wildtype (31.2±0.5 das; 21.4±1.2 leaves), *flac* single mutant plants bolt already slightly earlier (29.0±0.6 das; *P* = 0.022) and have fewer rosette leaves at this point (16.9±1.0 leaves; *P* = 0.036) (Fig 4A and B). In the *flic flac* double mutant, this early-flowering phenotype is strongly enhanced as bolting occurs already at 23.5±0.5 das (compared to the wildtype and *flac*: *P* < 0.001) and double mutant plants possess an average of 10.8±0.3 leaves (compared to the wildtype and *flac*: *P* < 0.001).

**Figure 4.**
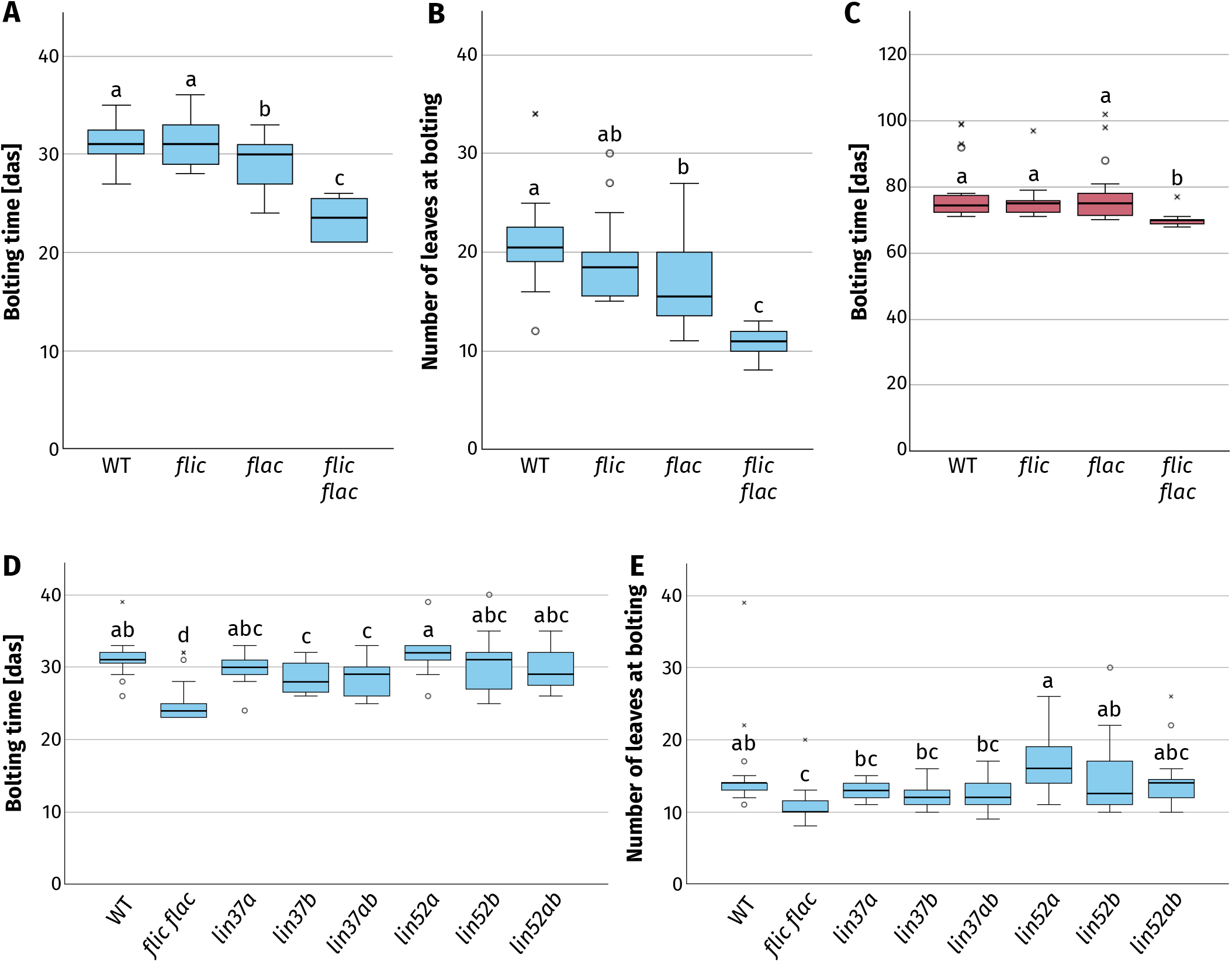
Flowering time analysis of *flic*, *flac*, *flic flac*, and their MuvB-core interactor (*lin37* and *lin52*) mutant plants. Floral transition was compared for mutants of *FLIC*, *FLAC*, and their MuvB-core interactors *LIN37* and *LIN52* by quantifying bolting as the time point when the shoot of a plant reached a height of 1 cm as well as the number of rosette leaves at that time point. Blue and pink boxes indicate LD and SD conditions, respectively. **(A)** Assessment of bolting time for wildtype, *flic*, *flac*, and *flic flac* mutants grown under LD conditions in days after sowing (das). **(B)** Number of rosette leaves at the time point of bolting for wildtype, *flic*, *flac*, and *flic flac* mutants grown under LD conditions. **(C)** Assessment of bolting time for wildtype, *flic*, *flac*, and *flic flac* mutants grown under SD conditions. **(D)** Assessment of bolting time for wildtype, *flic flac*, and their MuvB-core interactor (*lin37* and *lin52*) mutants grown under LD conditions. **(E)** Number of rosette leaves at the time point of bolting for wildtype, *flic flac*, and their MuvB-core interactor (*lin37* and *lin52*) mutants grown under LD conditions.

Next, we wanted to determine whether FLIC and FLAC control flowering in a photoperiod-dependent manner and measured the time to bolting in short-day (SD) conditions, for which plants were grown in an 8 h light/16 h dark cycle (Fig 4C). While we could not observe a significant early-flowering phenotype for the *flac* single mutant in these conditions (77.5±2.1 das), barely differing from the wildtype (78.3±2.1 das), the *flic flac* double mutant again showed significantly early floral initiation (69.8±0.4 das; *P* = 0.004). These results demonstrate that FLIC and FLAC are general regulators of flowering, independent of the light regime.

Since we could identify LIN37 and LIN52 as direct interactors of FLIC and FLAC, we assessed whether these MuvB-core components were also involved in flowering time control. To this end, we performed the same LD assay as before for all *lin37* and *lin52* single and double mutants (Fig 4D and E). Whereas *lin52* mutant (*lin52a-c1*: 31.8±0.7 das; 16.8±1.0 leaves; *lin52b-1*: 30.4±0.8 das; 14.8±1.1 leaves; *lin52ab*: 29.7±0.6 das; 14.3±0.8 leaves) and *lin37a* single mutant (29.7±0.4 das; 12.9±0.3 leaves) bolting times appeared to be similar to the wildtype (31.2±0.6 das; 15.2±1.4 leaves), we found a significant difference when abolishing *LIN37B*. Both *lin37b* single mutants (28.5±0.5 das, *P* = 0.048; 12.2±0.3 leaves) and *lin37ab* double mutants (28.5±0.6 das, *P* = 0.040; 12.5±0.4 leaves) were bolting early, however, not to the degree that we could observe for the *flic flac* double mutant (25.2±0.7 das, compared to *lin37b* and *lin37ab*: *P* < 0.01; 10.9±0.6 leaves).

These findings demonstrate that the DREAM complex not only plays a role in the promotion of floral transition as suggested previously, but is also repressing it via the MuvB-core component LIN37B.

### LIN37B appears to facilitate a ternary interaction connecting JMJ14/NAC050/NAC052 and FLIC/FLAC

Our above-presented finding that FLIC and FLAC are in one protein interaction network with the flowering time regulators JMJ14 and NAC052 suggested that they possibly directly interact to control flowering. To test this, we generated split-YFP fusion constructs to perform BiFC assays by transient tobacco transformation. Since a flowering time phenotype could be seen for *lin37* mutants, we decided to assess the interactions of FLIC, FLAC, and both LIN37 homologues each with JMJ14, NAC050, as well as NAC052 (Fig 5A). In our BiFC experiments, we were able to detect a YFP signal for almost all of the combinations tested (except for the three pairs of cYFP- FLAC/nYFP-NAC050, cYFP-LIN37A/nYFP-JMJ14, and cYFP-LIN37A/nYFP-NAC052).

**Figure 5.**
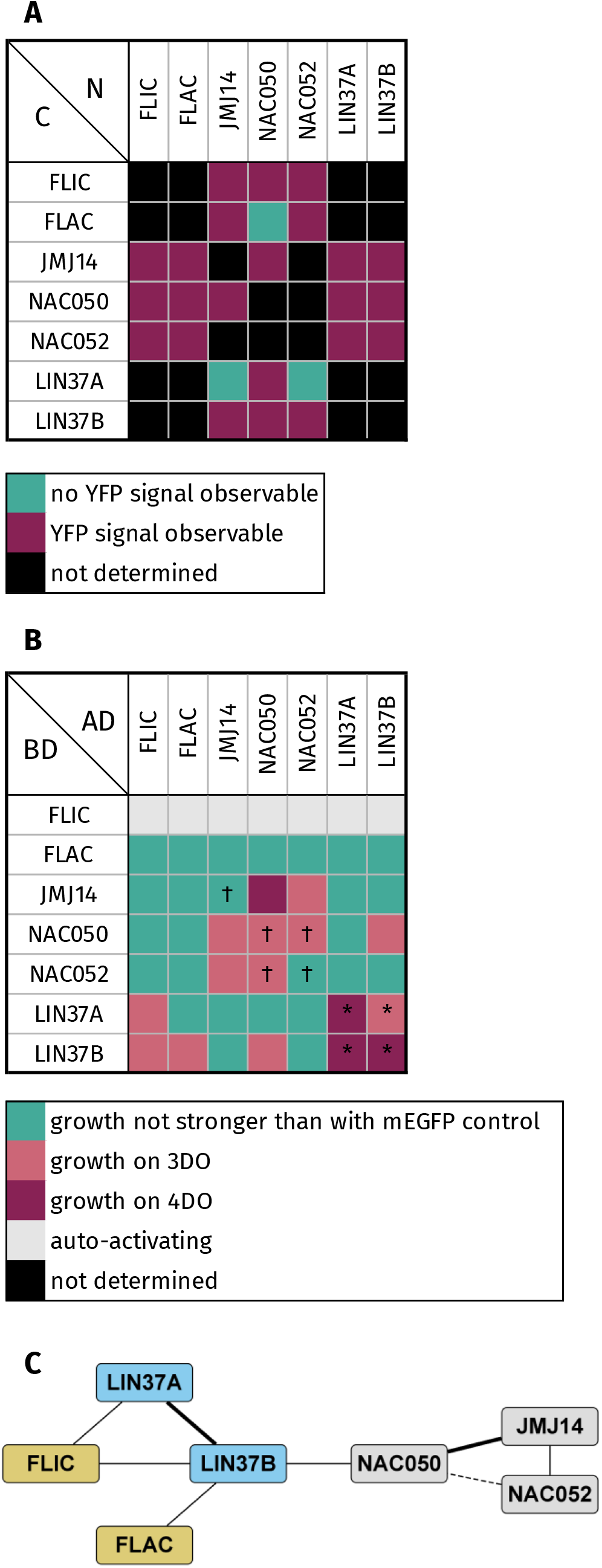
Interaction assays of FLIC, FLAC, LIN37 and the JMJ14/NAC050/NAC052 module. Interactions between FLIC, FLAC, their early-flowering MuvB-core interactors LIN37A and LIN37B, as well as JMJ14, NAC050, and NAC052 were determined via BiFC and Y2H assays. **(A)** BiFC interaction matrix. Cyan, no YFP signal could be detected; pink, YFP signal could be detected; black, not determined. **(B)** Y2H interaction matrix. Interactions of FLIC, FLAC, and LIN37 with JMJ14, NAC050, and NAC052 were tested with N- and C-terminal fusion proteins. Only the strongest signal obtained is shown for each combination. *, interaction determined previously (Lang et al, 2021). †, interaction determined previously (Ning et al, 2015). Dark pink, signal on 4DO; red, signal on 3DO but not on 4DO; cyan: signal not stronger than with mEGFP control; grey, strong auto-activation observable; black, not determined. **(C)** Schematic representation of the interactions observed in the Y2H assays (self-interactions were omitted). Thick line, signal on 4DO; thin solid line, signal on 3DO but not on 4DO; thin dashed line, as determined previously (Ning et al, 2015). Gold, FLIC/FLAC; blue, MuvB- core protein; dark cyan, other DREAM protein.

To corroborate our BiFC data, we next tested the same combinations in Y2H assays with N- and C-terminally tagged fusions. When fused to the BD, FLIC showed a strong auto-activating activity in both variants. We could determine that while FLIC, FLAC, and the JMJ14/NAC050/NAC052 module show no binary interactions, FLIC, FLAC, and NAC050 share LIN37B as a common interactor (Fig 5B and C). Taken together, our findings open up the possibility of a ternary interaction linking FLIC/FLAC and the JMJ14/NAC050/NAC052 module indirectly.

## Discussion

The repression of premature floral transition and its subsequent onset at the optimal time requires tight regulation to ensure maximal reproductive fitness. This intricate process is governed by multiple pathways influenced by a plethora of internal and external factors that collectively determine the timing of floral meristem identity establishment. Here, we demonstrate a role of the DREAM complex and its novel plant-specific components FLIC and FLAC in proper maintenance of floral repression likely working in conjunction with JMJ14, NAC050, and NAC052.

When performing AP-MS experiments with FLIC and FLAC as baits, we were able to co-purify a large part of the DREAM complex, including at least one homologue of each MuvB-core component (Fig 1, Tables S1 and S2), placing these proteins as novel plant DREAM components. However, whether and to what degree the mutant phenotypes observed for *flic* and *flac* are due to their function within a DREAM complex versus an independent role remains to be investigated.

Evidence for a repressive nature of a FLIC/FLAC-containing DREAM complex comes with the observation that both proteins interact with BTE1/BTL1. It has recently been published that BTE1 is required for transcriptional repression by the DREAM complex through binding of WD REPEAT DOMAIN 5A (WDR5A) and denying its binding to chromatin. Thereby, H3K4me2 deposition is facilitated, whereas H3K4me3 deposition is inhibited (Wang et al, 2022b). Interestingly, it has already been demonstrated that WDR5A is repressing the floral transition, with mutants exhibiting an early-flowering phenotype, by regulating methylation levels of the floral repressor genes *FLC* and *MAF4* (Jiang et al, 2009; Zhao et al, 2018). The recent identification of WDR5A in Co-IPs with TRB protein baits, while TRBs could be shown to interact with JMJ14, hints at the possible existence of a complex network (Fig 6) (Wang et al, 2023). Regarding the DREAM-related BTE1/BTL1 interactome besides FLIC and FLAC, we could establish that their only anchor point to the MuvB-core is LIN37B. Apart from this, BTE1 and BTL1 rather appear to interact with other cell cycle regulatory proteins of the DREAM complex as we could detect interactions with DPA, E2FE, and MYB3R1 (Fig S2).

**Figure 6.**
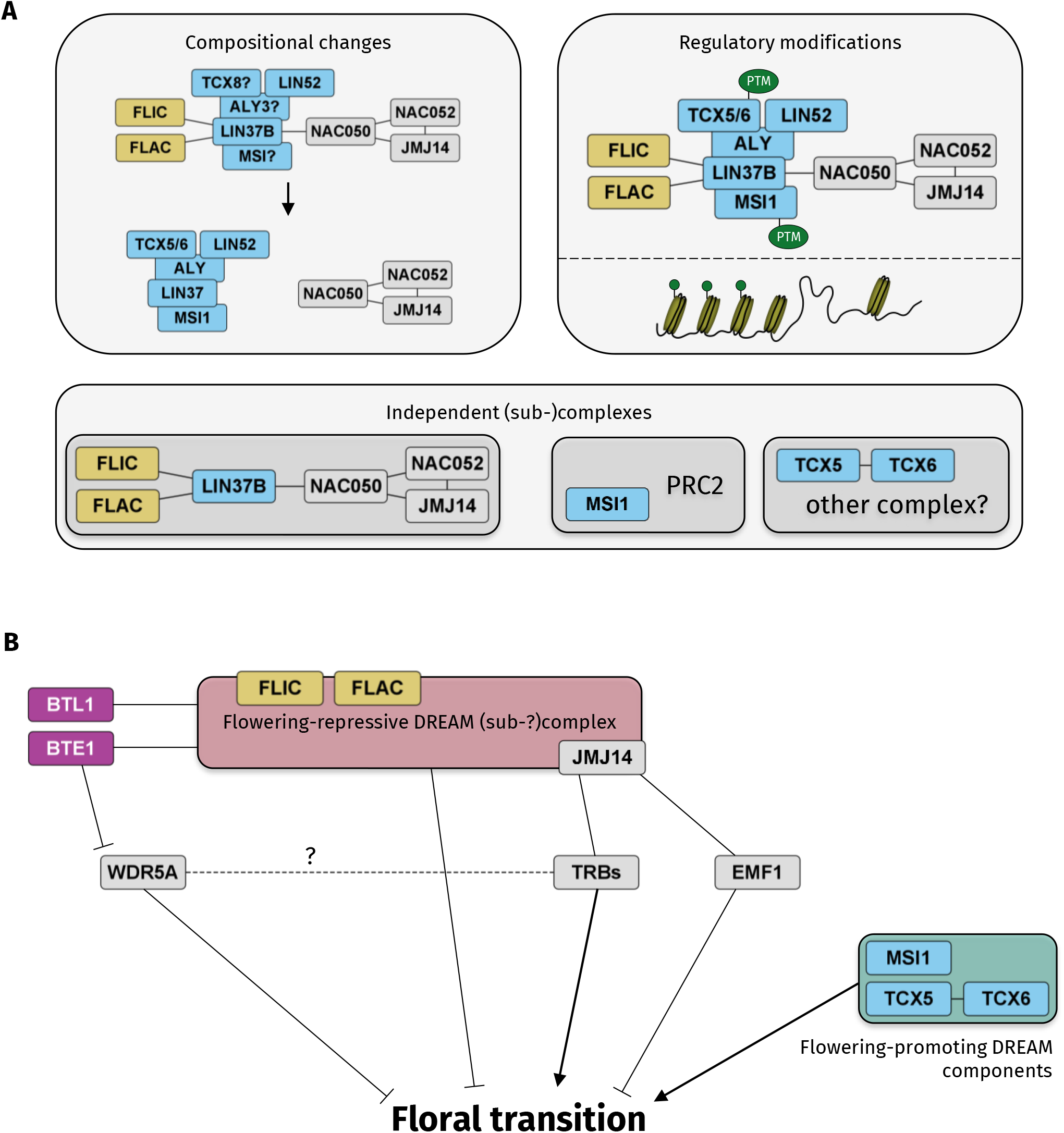
Model of DREAM involvement in flowering time regulation. **(A)** Possible modes of action of the DREAM complex to control flowering. Compositional changes could establish separate variants of the complex to either repress or promote flowering at different time points. Regulatory modifications, such as post-translational modifications (PTMs) or histone marks (green circles), might govern the ability of individual subunits to regulate transcription, even if they are part of the same complex. Lastly, DREAM components possibly form independent (sub-)complexes. Note, that the presented options are not mutually exclusive. **(B)** A variant or subcomplex of the DREAM complex (as presented in (A)) including FLIC, FLAC, LIN37B, and the JMJ14/NAC050/NAC052 module is repressing flowering when appropriate. Additionally, JMJ14 seems to be recruited by different transcription factors, both repressing and promoting flowering, possibly depending on environmental cues or developmental requirements. Other components of the DREAM complex, i.e. MSI1, TCX5, and TCX6, have been shown to promote flowering. BTE1 and BTL1, which are directly binding to components of the flowering-repressive complex, appear to indirectly promote floral transitioning by binding to and blocking of WDR5A action. WDR5A has recently been identified in Co-IPs using TRB baits indicating a possible connection, however if so, the nature of it is still unclear.

Interestingly, we also identified the histone demethylase JMJ14 in our pull- downs. Outside of the plant kingdom, JmjC domain-containing histone demethylases have been extensively characterised. Functions in a range of nuclear processes, such as transcriptional control, cell cycle transitions, and DNA repair have been shown, and they are involved in a diverse set of diseases, including cancer, cardiac disease, and obesity (Franci et al, 2014; Dimitrova et al, 2015). The roles of this class of proteins in plants are being progressively elucidated (Crevillén, 2020; He et al, 2021; Ma et al, 2022; Wang et al, 2022a).

Mutants for another JmjC domain-containing histone demethylase INCREASE IN BONSAI METHYLATION 1 (IBM1) exhibit developmental defects similar to *flic flac* mutants (Saze et al, 2008). Remarkably, it has been demonstrated that a mutation of the chromatin remodeler *DECREASED DNA METHYLATION 1* (*DDM1*), which in turn leads to hypomethylation, results in a similar dwarfism phenotype by the constitutive overexpression of pathogenesis-related (PR) genes. This happens via deregulation of the SA-dependent defence response pathway and is therefore striking the link to phytohormonal imbalance (Stokes et al, 2002; Li et al, 2010; Zhu et al, 2010). Further, DDM1 plays an important role in flowering time regulation via maintaining methylation levels of FWA (Kakutani, 1997; Soppe et al, 2000). As the deregulated methylation patterns in *ibm1* and *ddm1* mutants differ, an additive phenotype can be observed in *ddm1 ibm1* double mutants (Saze et al, 2008). Since the DREAM complex has recently been linked to methylation maintenance in several ways (Ning et al, 2020; Wang et al, 2022b), FLIC and FLAC might be involved in this process as well, and it will be interesting to explore this possibility in future.

JMJ14 itself has been shown to associate with a diverse set of partners to regulate floral transition, such as EMF1 (Wang et al, 2014), the NAC transcription factors NAC050 and NAC052, and most recently TRBs. In this study, we unveil a novel regulatory mechanism of flowering time involving the DREAM complex and in particular its subunits FLIC and FLAC, as well as the core subunit LIN37B as novel interactors of the JMJ14/NAC050/NAC052 module. Ultimately, we could demonstrate that FLAC alone and FLIC in combination with its homologue are indispensable for proper regulation of floral transition in LD and SD conditions (Fig 4). The early-flowering phenotype we observed is very reminiscent of what has been described for plants lacking JMJ14 or NAC052, both of which we could identify in our pull-downs. A milder dysregulation of flowering time could be seen in mutants of *LIN37B*, a shared MuvB-core interactor of FLIC, FLAC, and the JMJ14/NAC050/NAC052 module via NAC050. These observations strongly point towards a co-operative repression of floral transition independent of photoperiod by JMJ14/NAC050/NAC052 and FLIC/FLAC facilitated via the DREAM complex.

Noteworthily, the DREAM complex seems to fulfil opposing roles of activating and repressing floral initiation through its different subunits. Whereas FLIC/FLAC and LIN37B delay flowering until appropriate, TCX5/6 and MSI1 are required for the timely activation of this process. This can be explained by the observation of the DREAM complex being able to change its functionality depending on its bound subunits and interactors (Fischer & DeCaprio, 2015; Engeland, 2018). However, at least MSI1 seems to also control flowering in the context of being incorporated into another complex, the PRC2 (Bouveret et al, 2006). While no research has been done on how TCX5 and TCX6 are promoting floral transition, we could not establish any function of LIN52 in flowering control. So far, we could not determine any mutant phenotype in *lin52ab* double mutants, and, in contrast to humans (Guiley et al, 2015), LIN52 does not bind RBR1 in Arabidopsis (Lang et al, 2021). Moreover, in other organisms, LIN52 has the crucial function of switching the complex from its repressive to an activating state after being phosphorylated at its serine residue 28 (Litovchick et al, 2011). This serine is not conserved in plants (Fig S4). Therefore, it is interesting to consider that LIN52 might have lost its functionality in plants at least partially.

Our observations lead us to hypothesise about at least three different modes of DREAM action that could reconcile its dual function in flowering time regulation (Fig 6A). Firstly, the composition of the DREAM complex might change depending on whether flowering should be repressed or activated. This could be realised through incorporation of different component homologues with separate sets of target genes, e.g. whereas TCX5 and TCX6 promote flowering, TCX8 is able to bind to ALY3 and represses LIPOXYGENASE 2 (LOX2) expression (Noh et al, 2021). Interestingly, it has been demonstrated that LOX2 is up-regulated upon floral transition, and there are hints at a delayed-flowering phenotype in the corresponding mutants (Bañuelos et al, 2008; Wang et al, 2017). Thus, TCX5 and TCX6 might even have opposing roles to other TCX homologues, such as TCX8. Such homologue-specific behaviour could also be the case for other DREAM components.

Secondly, regulatory modifications could govern the ability of individual DREAM components to control transcription. This possibly includes post-translational modifications of distinct subunits as well as epigenetic marks on specific genes to regulate chromatin access. Lastly, flowering regulation by DREAM components might be achieved through different (largely) independent (sub-)complexes. Just as MSI1 works in conjunction with the PRC2, LIN37 might act in an independent FLIC/FLAC–LIN37B–JMJ14/NAC050/NAC052 complex. Of course, it is possible that these regulational options overlap and apply at the same time. Further research will elucidate the true nature of DREAM action on flowering initiation and to what degree FLIC and FLAC are involved in this process.

Taking into account what has been discussed, it seems possible that these regulatory complexes extend to include BTE1/BTL1, WDR5A, and/or TRB proteins and that JMJ14 is recruited to multiple subsets of genes by different transcription factor interactors (Fig 6B). Thus, we are still beginning to unravel the stunning complexity of DREAM functionality in plants. It is already clear that the composition of the complex and the cross-talk with other complexes, such as the PRC2, will be decisive for its action in different developmental and environmental contexts.

## Material and Methods

### AP-MS and filtering of the data

Cloning of FLIC and FLAC encoding C- and N-terminal GS^rhino^ tag (Van Leene et al, 2015) fusions under control of the constitutive cauliflower tobacco mosaic virus 35S promoter and transformation of Arabidopsis cell suspension cultures (PSB-D) with direct selection in liquid medium was carried out as previously described (Van Leene et al, 2022).

Pull-downs were performed in triplicate, using in-house prepared magnetic IgG beads and 25 mg of total protein extract per pull-down as described (Van Leene et al, 2022). On-bead digested samples were analysed on a Q Exactive (ThermoFisher Scientific) and co-purified proteins were identified with Mascot (Matrix Science) using standard procedures (Van Leene et al, 2022).

After identification, the protein list was filtered versus a large dataset of similar experiments with non-related baits using calculated average Normalised Spectral Abundance Factors (NSAFs) (Van Leene et al, 2022). Proteins identified with at least two matched high confident peptides in at least two experiments, showing high and significant enrichment compared to the large dataset, either at least a 10- fold enrichment with a −log10(*p*-value(T-test))≥10, or at least 20-fold enrichment with a −log10(*p*-value(T-test))≥8, were retained. Furthermore, proteins appearing with only very few bait groups in the large dataset were retained when they met the following criteria: present with not more than two bait groups in the large dataset, and identified with at least two matched high confident peptide sequences in at least two experiments. Or present with no more than four bait groups in the large dataset and identified with one high confident peptide sequence in all three replicates and showing high (at least 10-fold) and significant [−log10(*p*-value(T-test))≥10] enrichment compared to the large dataset.

### Multiple sequence alignments

Alignments were generated using the MUSCLE algorithm using default settings (Edgar, 2004). Structural predictions were generated by AlphaFold (Varadi et al, 2022). Data visualisation was done using ESPript 3 (Robert & Gouet, 2014).

### Yeast two-hybrid assay

To generate constructs for yeast two-hybrid assays, the coding sequences of the respective genes were amplified from cDNA and attB-recombination sites were added in two consecutive PCRs. By Gateway BP reactions, these sequences were subcloned into the *pDONR223* entry vector. The corresponding N-terminally fused *pGAD424-GW* and *pGBT9-GW* expression clones as well as the C-terminally fused *pGADCg* and *pGBKCg* expression clones were generated by Gateway LR reactions. Primers used for construct generation are shown in Table S2.

Yeast two-hybrid assays were performed according to the Yeastmaker Yeast Transformation System 2 manual (Clontech). The yeast strain AH109 was co-transformed with an AD-fused and a BD-fused construct using the lithium acetate method. Yeast cells harbouring both constructs were grown on 2DO, 3DO, and 4DO medium (–L/–W, –L/–W/–H, and –L/–W/–H/–Ade, respectively) to assess protein/protein interactions. Co-transformation of a construct with the corresponding mEGFP construct was used as an auto-activation control.

### Plant materials and growth conditions

The *Arabidopsis thaliana* accession Columbia-0 (Col-0) was used as the wild-type reference. All mutants and transgenic lines used in this study were in the Col-0 background. The mutant lines *flic-2* (SAIL_893_B04), *flic-4* (GK-612E12), and *flac-1* (GK- 327D05) were identified from the GABI-KAT (Kleinboelting et al, 2012) or SAIL (Sessions et al, 2002) T-DNA collections and provided by the Nottingham Arabidopsis Stock Centre (NASC) (Scholl et al, 2000). The mutant lines *lin37a-2*, *lin37b-3*, *lin52a-c1*, and *lin52b-1* have been described previously (Lang et al, 2021).

Plants were germinated and grown on vertical plates containing ½ Murashige and Skoog (MS) medium under long-day conditions (16 h light, 8 h dark) at 22°C for 7 d. Seedlings were then transplanted into soil and grown under the same conditions for all experiments using adult plants, and additionally under short-day conditions (8 h light, 16 h dark) for the examination of bolting.

### Bimolecular fluorescence complementation assay

To generate constructs for bimolecular fluorescence complementation assays, the coding sequences of the respective genes were amplified from cDNA and attB-recombination sites were added in two consecutive PCRs. By Gateway BP reactions, these sequences were subcloned into *pDONR221-P1P4* and *pDONR221-P3P2* entry vectors. The corresponding *pBiFC-2in1-NN* expression clones were generated by Gateway LR reactions. Primers used for construct generation are shown in Table S2. The relevant proteins were transiently expressed in *Nicotiana benthamiana* leaves after *Agrobacterium tumefaciens* infiltration and the fluorescence of YFP was imaged 2 d after infiltration using a Leica SP8 laser-scanning confocal microscope.

### Pollen and seed viability assays

The Peterson staining method was used to analyse the pollen viability (Peterson et al, 2010). For counting of pollen, three mature flower buds containing dehiscent anthers (or five in the case of *flic flac*) were collected and dipped in 13 μL Peterson staining solution (25% glycerol, 10% ethanol, 4% glacial acetic acid, 0.05% acid fuchsin, 0.01% malachite green, 0.005% orange G) for 10 s on a microscope slide, which was then covered by a coverslip. Subsequently, slides were heated on a hotplate at 80°C for 10 min to distinguish aborted and non-aborted pollen grains. Slides were analysed and imaged using a light microscope. Seed sets were determined by quantifying viable and aborted seeds of mature siliques; 5 siliques of 5 plants per genotype were analysed.

## Data Availability

Supporting data for Fig 1 (AP-MS results) can be found in Tables S1 and S2.

## Acknowledgements

We are grateful to Masaki Ito (School of Biological Science and Technology, Kanazawa University) and Claudio Alfieri (The Institute of Cancer Research, London) for critical reading and helpful comments on the manuscript. We thank the VIB Proteomics Core Facility (VIB, University of Gent, Belgium) for performing the LC-MS/MS analyses. This work was supported through a fellowship of the University of Hamburg to LL, a fellowship of the Friedrich-Ebert-Stiftung to FB, and a DFG grant (SCHN 736/16-1) to AS.

## Author contributions

L Lang: conceptualisation, formal analysis, supervision, visualisation, writing— original draft.

F Böwer: formal analysis, investigation, and visualisation.

H Tunçay Elbaşı: investigation and visualisation.

D Eeckhout: data curation, formal analysis, and investigation. N Marschlich: investigation and visualisation.

G De Jaeger: resources, data curation, supervision, funding acquisition, and project administration.

M Heese: conceptualisation and formal analysis.

A Schnittger: conceptualisation, formal analysis, supervision, funding acquisition, writing—original draft, and project administration.

## Supplementary Figures

**Figure S1.**
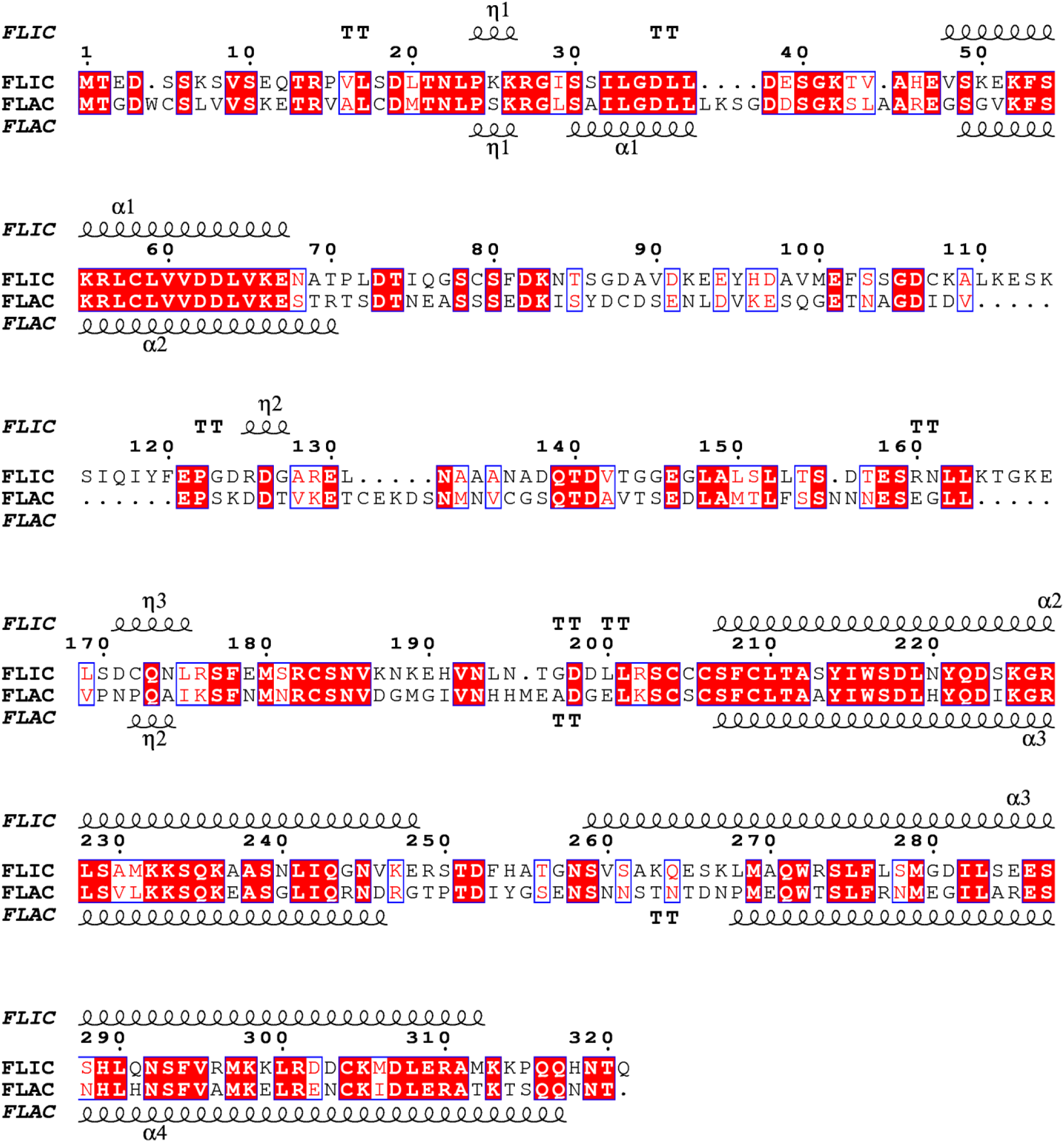
Pairwise sequence alignment of FLIC and FLAC protein sequences. Structural predictions for FLIC and FLAC are indicated above and below the sequences, respectively.

**Figure S2.**
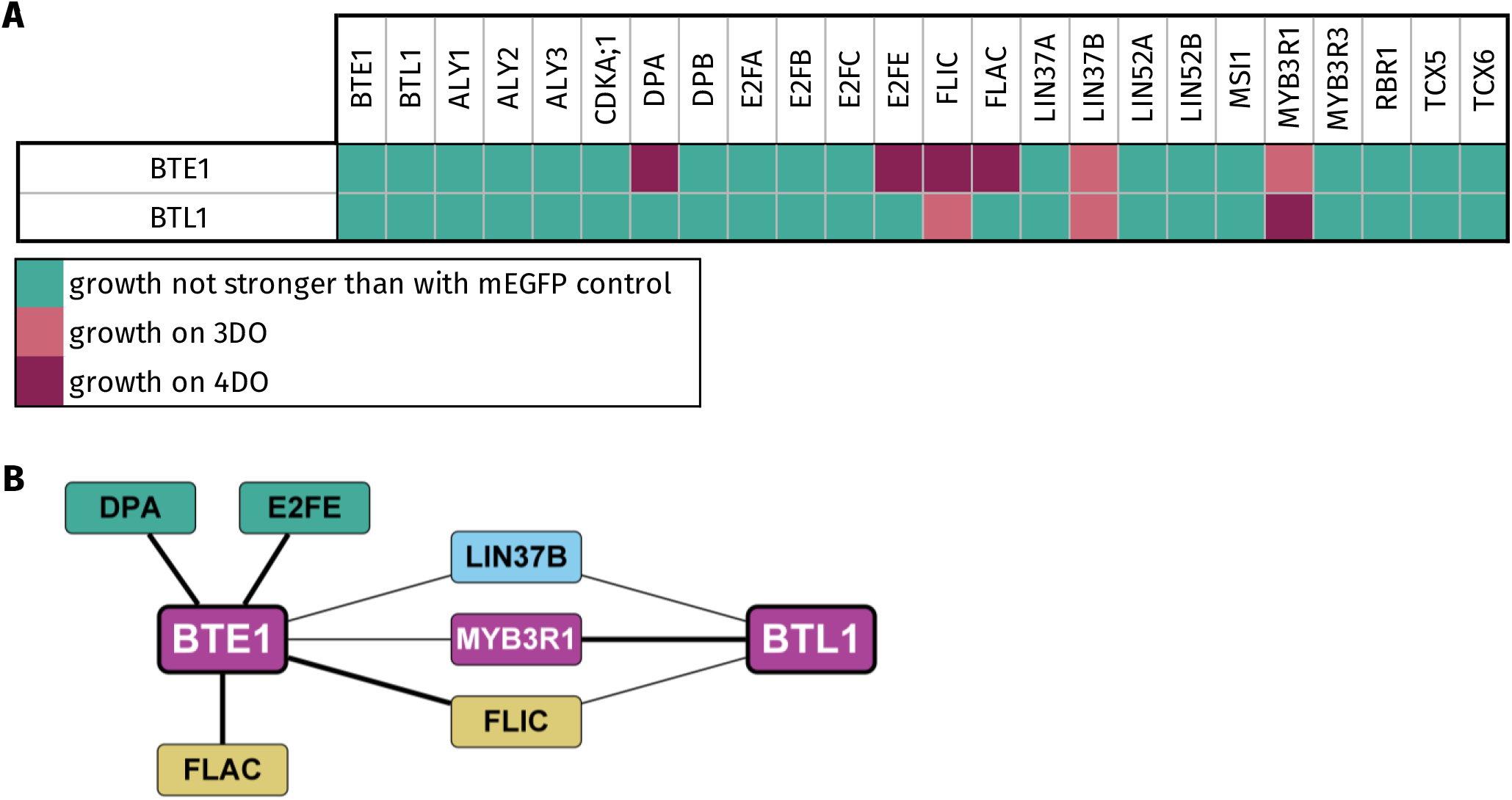
Binary interactions of BTE1 and BTL1 with the DREAM complex and associated proteins by Y2H. All proteins were tested using both AD- and BD-variants and only the strongest interaction is shown. Signal strength was classified according to yeast growth on different dropout media in three categories. **(A)** Dark pink, growth on 4DO; red, growth on 3DO but not on 4DO; cyan: growth not stronger than with mEGFP control. **(B)** Schematic representation of the observed interactions of BTE1 and BTL1 as indicated by edges. Thick line, growth on 4DO; thin line, growth on 3DO but not on 4DO. Gold, FLIC/FLAC; blue, MuvB-core protein; dark cyan, other DREAM protein; pink, DREAM-related proteins.

**Figure S3.**
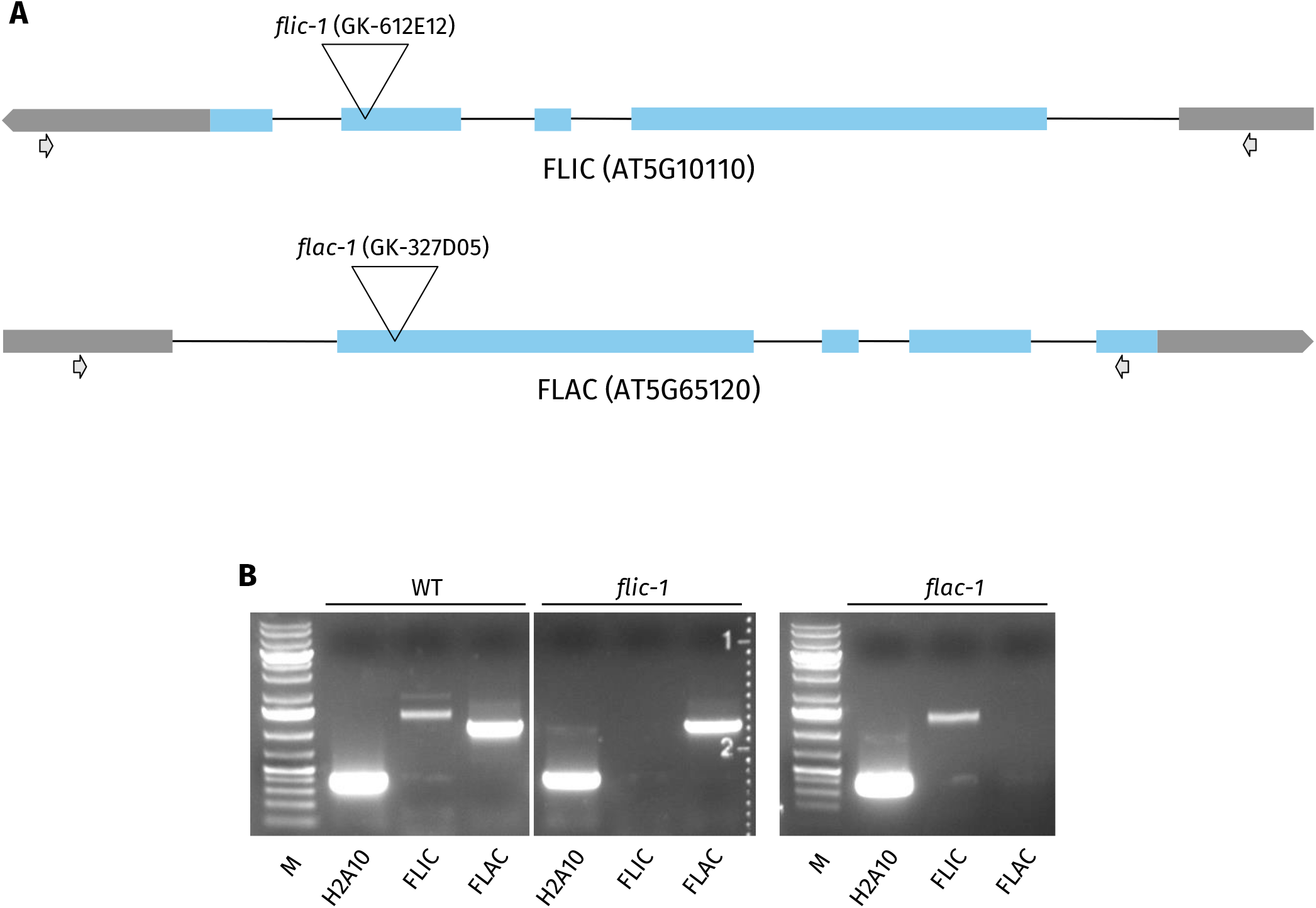
Scheme of the *FLIC* and *FLAC* genes and Arabidopsis mutants isolated in this study. T-DNA insertion positions were confirmed by sequencing of PCR products. The line GK-612E12 (*flic-1*) has its T-DNA insertion in the third exon of *FLIC*. An insertion line for *FLAC*, GK-327D05 (*flac-1*), shows an insertion in the first exon. **(A)** The T-DNA insertion sites of the mutant alleles as well as the location of primers used for expression analysis are indicated. Blue rectangles indicate exons, whereas black lines in between the rectangles show introns. Grey rectangles and pentagon arrows represent 5′ UTRs and 3′ UTRs, respectively. **(B)** The presence of full-length transcripts in the different insertion lines was checked by RT-PCR using H2A10 as a control for the generated cDNA. WT, Wildtype; M, GeneRuler DNA Ladder Mix.

**Figure S4.**
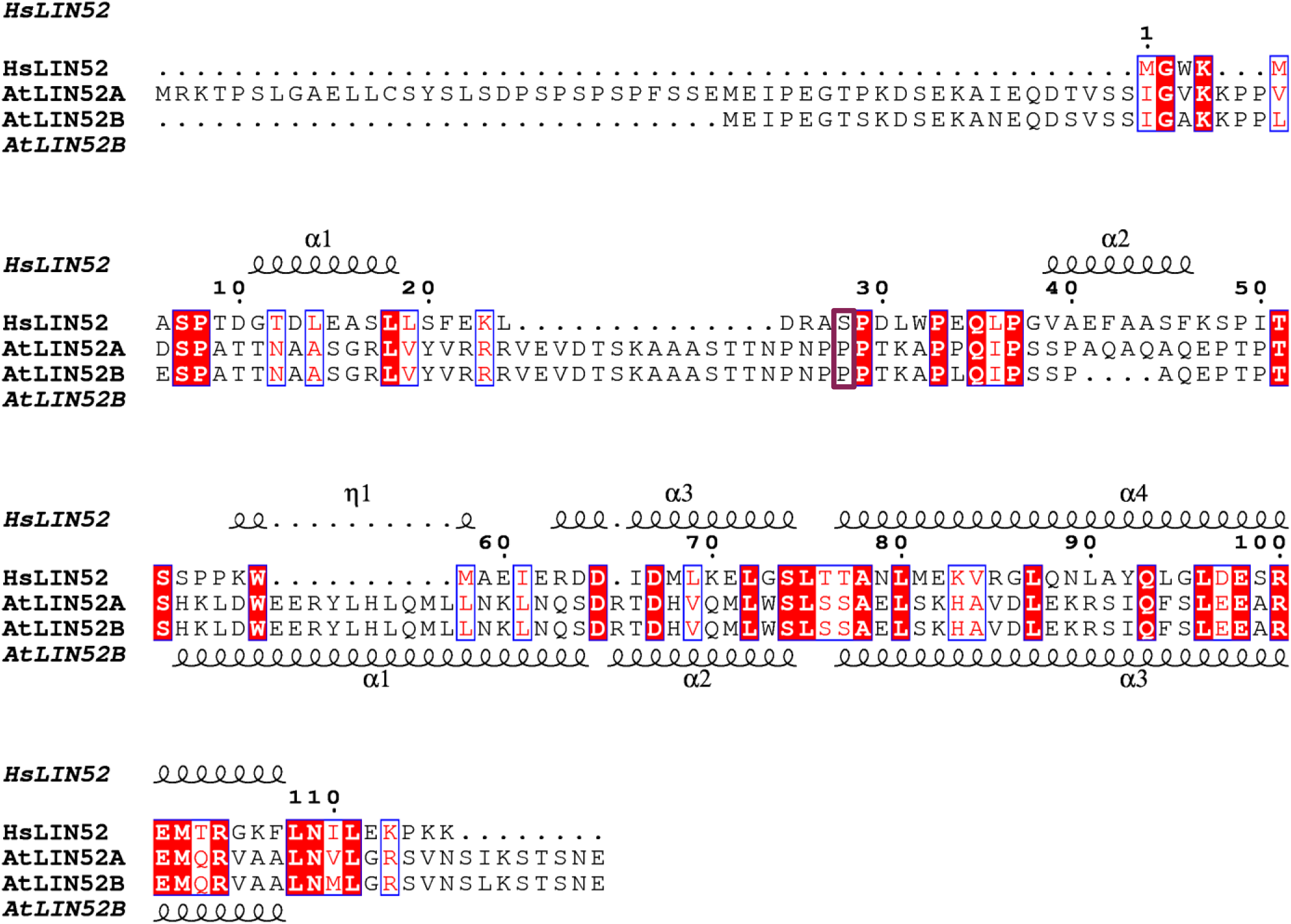
Multiple sequence alignment of HsLIN52 and Arabidopsis orthologue protein sequences. Structural predictions for HsLIN52 and AtLIN52B are indicated above and below the sequences, respectively. HsLIN52’s serine residue 28 and its proline substitutions in the Arabidopsis orthologues are surrounded by a red rectangle.

## Supplementary Tables

**Table S1.**

Overview of AP-MS results using FLIC and FLAC as baits (for details refer to Table S2). Numbers in columns C–F indicate the number of times the interactor was identified above threshold in the corresponding AP-MS replicates; only interactors that were found in at least 2 replicates of one experiment are considered reliable positives and are listed here.

**Table S2.**

Protein Identification details obtained with the Q Exactive (Thermo Fisher Scientific) and Mascot Distiller software (version 2.5.0, Matrix Science) combined with the Mascot search engine (version 2.6.2, Matrix Science) using the Mascot Daemon interface and database Araport11plus_DE2020 (database available from Pride repository project PXD029833) (Van Leene et al, 2022).

**Table S3.**

List of primers used in this study.

